# Acute SARS-CoV-2 infection is associated with an expansion of bacteria pathogens in the nose including *Pseudomonas Aeruginosa*

**DOI:** 10.1101/2021.05.20.445008

**Authors:** Nicholas S. Rhoades, Amanda Pinski, Alisha N. Monsibais, Allen Jankeel, Brianna M. Doratt, Isaac R. Cinco, Izabela Ibraim, Ilhem Messaoudi

## Abstract

Much of the research conducted on SARS-CoV-2 and COVID-19 has focused on the systemic host response, especially that generated by severely ill patients. Very few studies have investigated the impact of acute SARS-CoV-2 within the nasopharynx, the site of initial infection and viral replication. In this study we profiled changes in the nasal microbial communities as well as in host transcriptional profile during acute SARS-CoV-2 infection using 16S amplicon sequencing and RNA sequencing. These analyses were coupled to viral genome sequencing. Our microbiome analysis revealed that the nasal microbiome of COVID patients was unique and was marked by an expansion of bacterial pathogens. Some of these microbes (i.e. *Acinetobacter*) were shared with COVID negative health care providers from the same medical center but absent in COVID negative outpatients seeking care at the same institutions suggesting acquisition of nosocomial respiratory pathogens. Specifically, we report a distinct increase in the prevalence and abundance of the pathogen *Pseudomonas aeruginosa* in COVID patients that correlated with viral RNA load. These data suggest that the inflammatory environment caused by SARS-CoV-2 infection and potentially exposure to the hospital environment leads to an expansion of bacterial pathogens in the nasal cavity that could contribute to increased incidence of secondary bacterial infections. Additionally, we observed a robust host transcriptional response in the nasal epithelia of COVID patients, indicative of an antiviral innate immune repones and neuronal damage. Finally, analysis of viral genomes did not reveal an association between viral loads and viral sequences.

## INTRODUCTION

Severe acute respiratory syndrome coronavirus 2 (SARS-CoV-2) is the causative agent of coronavirus disease 2019 (COVID-19) and the cause of the ongoing global pandemic(*1, 2*). COVID-19 is a respiratory disease ranging in clinical presentation from an asymptomatic (~80% of cases) to a fatal infection (~1-2%) accompanied by cytokine storm, acute respiratory distress syndrome and coagulopathy (*3, 4*). Long-term consequences of COVID-19 range from anosmia and ageusia(*3, 5, 6*) to respiratory (e.g., dyspnea) and nervous system (e.g., headache, anosmia) complications (*7*). Numerous studies have identified advanced age, comorbidities (notably, obesity and diabetes), and biological sex as risk factors for severe COVID-19 (*4, 5, 8, 9*). Other factors modulating disease, including mucosal microbial defenses and viral burden at the site of pathogenesis have been under-explored.

SARS-CoV-2 is primarily spread through the inhalation of virus-laden respiratory droplets(*5*). Initial replication occurs in the upper respiratory tract (URT) following the interaction of the viral spike protein and host ACE2, and subsequent entry into cells (*10–12*). The nasal cavity is believed to be the initial site of viral replication rather than the oral cavity due to higher expression of ACE2 (*13–15*) and greater levels of viral RNA (vRNA) in this site compared to matched oral specimens (*16*). Rapid and uncontrolled replication can lead to infection of the lower respiratory tract (LRT) and severe disease. However, several studies have demonstrated a lack of correlation between disease severity and viral RNA load in nasal cavities, suggesting that other factors contribute to disease at the site of infection (*17–19*).

Recent research has demonstrated an increasingly critical role of the mucosal microbiome in modulating viral infection (*20–23*). Disruption of microbial communities by viral infection can exacerbate inflammation, facilitate coinfection, and deregulate the adaptive immune response (*20–23*). Furthermore, severe inflammation in olfactory epithelium can lead to short- and long-term symptoms like anosmia and ageusia(*24, 25*). However, while numerous studies have investigated the longitudinal dynamics of COVID-19 in blood, a limited number of studies have examined the respiratory microbiome in acute patients(*26–31*). Studies of nasopharyngeal samples are limited to small cohorts of mild and severe COVID-19 patients and show variable differences in diversity and increased abundance of select phyla without accounting for the potential impact of viral burden(*32–35*). Other studies of bronchoalveolar lavage and pharyngeal samples in severe patients have identified a general dysbiosis of the lower respiratory microbiome similar to that of pneumonia patients(*36, 37*).

Studies aimed at understanding the impact of COVID-19 and viral burden on nasal microbial communities is clearly lacking but urgently needed to characterize both acute infection and provide insight into long-term symptoms of COVID-19. In this study, we determined the impact of acute SARS-CoV-2 infection with a spectrum of viral RNA loads on nasal microbial communities, local host transcriptional responses and potential association with viral genome sequence diversity. Our cross-sectional study of hospitalized COVID-19 patients, and uninfected outpatients and healthcare workers indicated a distinct shift in the nasal microbiome of COVID-19 patients featuring the expansion of pathobionts such as *Rothia*, *Acinetobacter*, and *Pseudomonas*. This was accompanied by the upregulation of host antiviral immune genes and the downregulation of genes with critical roles in mucosal and neuronal cell homeostasis. However, no association between viral loads and viral genome sequences was detected. Overall, our data suggests SARS-CoV-2 infection is associated with an expansion of bacterial pathogens in the nasal cavity and virus-induced host dysregulation at the site of infection.

## METHODS

### Sample collection and extraction

RNA was extracted from remnant de-identified nasal swab samples collected in viral transport medium (VTM) from patients and healthcare workers (HCW) at the University of California, Irvine Medical Center (UCIMC) using the Quick-RNA™ Viral 96 Kit (Zymo, Cat#R1041). SARS-CoV-2 viral loads were determined using qPCR with primers specific to the nucleoprotein (NP). A one-step reaction was prepared using 5ul of extracted RNA or standard, 500nM of forward (5’-GGGGAACTTCTCCTGCTAGAAT-3’) and reverse (5’-CAGACATTTTGCTCTCAAGCTG-3’) SARS-CoV-2-nucleocapsid primers, 125 nM of SARS2-nucleocapsid probe (5’-FAM-TTGCTGCTGCTTGACAGATT-BHQ1-3’), and TaqPath™ 1-Step RT-qPCR Master Mix (Applied Biosystems, Foster City, CA, USA). PCR cycler conditions were 2 min at 95 °C, 15 min at 50 °C, denaturation for 2 min at 95 °C followed by 45 cycles of 3 s at 95 °C and 30 s 60 °C on the StepOnePlus™ Real-Time PCR System (Applied Biosystems, Foster City, CA, USA). All samples were run in triplicate. Viral load concentration for the target samples was determined by plotting the average Ct values against the log concentration of the standards in the dilution series. Positivity was determined as a cycle threshold (Ct) value less than or equal to 40 for the RBD gene. Samples were categorized into three groups: healthcare workers (HCW; n=45), patients testing negative for SARS-CoV-2 (CoV-; n=21), and patients who tested positive for SARS-CoV-2 (CoV+; n=68). The latter group was further sub-divided into three groups based on Ct values: High Ct value (>34.6; n=23), Mid Ct value (32.9-25.0; n=23), and Low Ct values (<23.5; n=22).

### 16S amplicon libraries construction and data analysis

DNA that co-eluted with extracted RNA was used as the template to amplify the hypervariable V4 region of the 16S rRNA gene using PCR primers (515F/806R with the forward primer containing a 12-bp barcode) in duplicate reactions containing: 12.5 ul GoTaq master mix, 9.5 ul nuclease-free H20, 1 ul template DNA, and 1 ul 10uM primer mix. Thermal cycling parameters were 94°C for 3 minutes; 35 cycles of 94°C for 45 seconds, 50°C for 1 minute, and 72°C for 1 minute and 30 seconds; followed by 72°C for 10 minutes. PCR products were purified using a MinElute 96 UF PCR Purification Kit (Qiagen, Valencia, CA, USA). Libraries were sequenced (2 x 300 bases) using Illumina MiSeq.

Raw FASTQ 16S rRNA gene amplicon sequences were uploaded and processed using the QIIME 2 version 2019.10 (*38*) analysis pipeline as we have previously described (*39*). Briefly, sequences were demultiplexed and quality filtered using the DADA2 plugin for QIIME 2 (*40*), which filters chimeric sequences. The generated sequence variants were then aligned using MAFFT (*41*), and a phylogenetic tree was constructed using FastTree 2 (*42*). Taxonomy was assigned to sequence variants using q2-feature-classifier against the SILVA Database (release: 138) (*43*). After taxonomic classification all sequences not assigned to a known phyla were removed along with any sequence assigned to mitochondria or chloroplast. After removal of low quality and contaminate sequencies taxonomy was reassigned to sequence variants using q2-feature-classifier against the expanded Human Oral Microbiome Database (eHOMD: release version 15.21) (*44*). This curated and site specific database was used to improve species level resolution. To prevent sequencing depth bias, samples were rarified to 8,000 sequences per sample before α and β diversity analysis. QIIME 2 was also used to generate the following α diversity metrics: richness (as observed taxonomic units), Shannon evenness, and phylogenetic diversity. β diversity was estimated in QIIME 2 using weighted and unweighted UniFrac distances (*45*).

All statistical analyses were conducted using PRISM (V8) or the R package Vegan (*46*). PERMANOVAs were performed using the Vegan function ADONIS. 1-way, non-parametric Kruskal-Wallis ANOVA were implemented using PRISM to generate p-values and utilizing the corresponding post-hoc-test when the initial ANOVA was significant. The LEfSe algorithm was used to identify differentially abundant taxa and pathways between groups with a logarithmic Linear discriminant analysis (LDA) score cutoff of 2 (*47*).

### RNAseq library prep and analysis

Quantity and quality of RNA extracted from 8 nasal swabs of (4 CoV+ and 4 HCW) was determined using an Agilent 2100 BioAnalyzer. cDNA libraries were constructed using the NEB Next Ultra II Directional RNA Library kit (Thermo Fischer). Briefly, RNA was treated with RNAseH and DNase I after depletion of ribosomal rRNA. Adapters were ligated to cDNA products. The ~300 bp amplicons were PCR-amplified and indexed with a unique molecular identifier. cDNA libraries were assessed for quality and quantity on the BioAnalyzer prior to single-end sequencing (x100bp) using the Illumina NovaSeq platform.

Bioinformatic analysis was performed using the systemPipeR RNA-Seq workflow (*48*). RNA-Seq reads were demultiplexed, quality-filtered and trimmed for quality and adapter removal using Trim Galore (average Phred Score cut-off = 30, minimum length of 50 basepairs). Tophat was used to align filtered reads to the reference genome *Homo sapiens* (GRCh38). The file was used for annotation. Raw expression values (gene-level read counts) were generated using the summarizeOverlaps function and normalized (read per kilobase of transcript per million mapped reads, rpkm) using the edgeR package. Statistical analysis with edgeR was used to determine differentially expressed genes (DEGs) meeting the following criteria: genes with median rpkm of ≥1, a false discovery rate (FDR) corrected p-value ≤ 0.05 and a log2fold change ≥ 1 compared to CoV-samples. Functional enrichment of DEGs was performed using Metascape to identify relevant Gene Ontology (GO) biological process terms (*49*). Heatmaps, bar graphs and volcano plots were generated using R package ggplot2.

### Viral genome library construction and analysis

Enrichment for viral RNA was performed using the QIASeq™ SARS-CoV-2 Primer Panel V2 (Qiagen) Panel V2) followed by library construction with the Qiagen FX DNA library preparation kit. Samples were indexed, pooled and validated with 2100 Agilent BioAn. Prior to Illumina sequencing (2×100bp, 1 M reads).

Adapters and primers were removed from demultiplexed sequencing readings using TrimGalore and MaskPrimers.py (pRESTO). Merged reads were aligned to SARS-CoV-2 Wuhan isolate NC_045512.2 with BWA-mem software version 0.7.17. Genomes with greater than 90% coverage and 10X depth were retained for phylogenetic analysis with Nextstrain. Amino acid changes and clade assignments were identified using Nextclade. A set of comparator sequences representing the original Wuhan isolate (NC_045512.2), variants of concern (B.1.351, EPI_ISL_960123; P.1, EPI_ISL_833167; B.1.1.7, EPI_ISL_659057), isolates from January 2020 New York (EPI_ISL_1293138) and Washington (EPI_ISL_404895), and representative isolates from California January-November 2020 (EPI_ISL_406036, EPI_ISL_411954, EPI_ISL_429875, EPI_ISL_436642, EPI_ISL_444023, EPI_ISL_569672, EPI_ISL_548382, EPI_ISL_548612, EPI_ISL_582897, EPI_ISL_625601, EPI_ISL_753324) were used for phylogenetic analysis. All sequences from Orange County, California from January 2020 to December 2020 were also retrieved from GISAID.

## RESULTS

### SARS-CoV-2 infection is associated with expansion of pathobionts in the nasal microbiome

To determine if SARS-CoV-2 infection is associated with a shift in the composition of the nasal microbiome, we utilized 16S rRNA gene amplicon sequencing to profile nasal samples from hospitalized COVID-19 patients (CoV+ inpatient), and SARS-CoV-2 negative out-patients (CoV-outpatient) and healthcare workers (CoV- HCW) (**Figure1A**). At the community level, the nasal microbiome of CoV+ patients was distinct from both CoV- HCW and CoV- individuals with host status accounting for 9.3% of the total variation of unweighted Unifrac distance (**Figure 1B**). Additionally, nasal microbial communities in CoV+ inpatients showed the highest intra-group variability followed by communities from CoV- HCW and CoV- outpatients (**Figure 1C**). These same trends held for abundance based weighted Unifrac distance (**Supp Figure 1A, B**). We also observed an increased richness in nasal community from COV+ patients and HCW as indicated by a higher number of observed Amplicon Sequencing Variants (ASVs) compared to CoV- outpatients (**Figure 1D**). As expected, all three groups had similarly low community Shannon diversity values, indicative of a low-complexity community dominated by a few microbes (**Figure 1E**). We next probed the taxonomic landscape of the nasal microbiome to identify microbial taxa that were driving the shift in the overall community. At the phyla level, samples from all groups were dominated by Actinobacteria, Firmicutes, and Proteobacteria (**Figure 1F, Supp. Figure 1C)**. Common nasal genera such as *Corynebacterium*, *Staphylococcus*, *Streptococcus*, *Dososigranulum*, and *Neisseria* were highly abundant in all three groups (**Figure 1F, Supp Figure 1C**).

**Figure 1:**
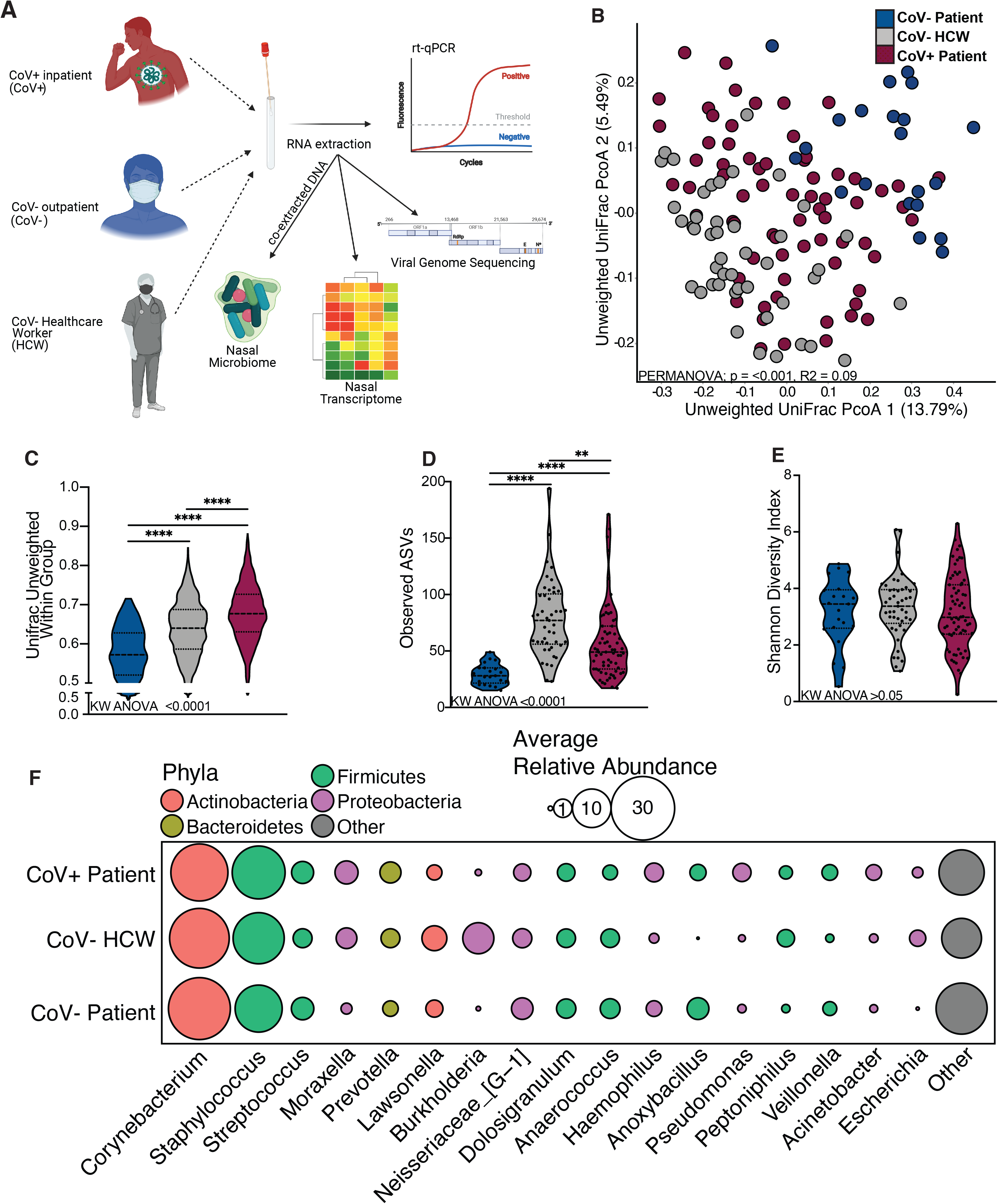
The nasal microbiome of SARS-CoV-2 infected patients is distinct. (A) Study design schematic. (B) Principal-coordinate analysis of nasal microbial communitues unweighted UniFrac distance colored by host status. The contribution of host status the total variance in the unweighted UniFrac dissimilarity matrices was measured using PERMANOVA (Adonis with 10,000 permutations). (C, D, E) Violin plot illustrating (D) average unweighted UniFrac distances (C) number of observed amplicon sequencing variants and (D) shannon diversity (E) split by host status. Significance for panels C-E was determined using Kruskal Wallis non-parametric ANOVA (p-values inset at the bottom of each panel), with Dunn’s multiple comparison * = p < 0.05, ** = p <0.01, *** = p < 0.001, **** = p < 0.0001. (F) Bubble plots of bacterial genera found at greater than 1% average abundance across the entire study population ordered from left to right in descending average abundance. The size of each circle indicates the average relative abundance for each taxa in the denoted group and the color of each circle denotes which bacterial Phyla each genera belongs to.

We next explored differences between the three groups. Analysis using LefSe revealed enrichment of specific genera in each group (**Figure 2A**, **Supp Table 1**). Specifically, *Anaerococcus* was enriched in CoV- outpatients, while *Lawsonella* and *Burkholderia* were most abundant in CoV- HCW (**Figure 2A**, **Supp Table 1**). Finally, several pathogenic bacteria such as *Rothia, Acinetobacter,* and *Pseudomonas* were most abundant in COV+ in-patients (**Figure 2A**, **Supp Table 1**). To further explore the distribution of specific taxonomic groups, we plotted the abundance of select taxa across all three groups. The two most abundant genera, *Corynebacterium* and *Staphylococcus* were equally abundant in the groups (**Figure 2B,C**). Interestingly, relative abundance of *Anoxybacillus* was higher in both outpatients CoV- and CoV+ inpatients compared to HCW (**Figure 2D**). Conversely, *Burkholderia cepacia* was significantly more abundant in CoV- HCW (**Figure 1F, 2E)**. On the other hand, the genera *Acinetobacter,* and *Pseudomonas* were most abundant in the COV+ inpatients (**Figure 2 F, G**). When probed at the species level, *Pseudomonas aeruginosa* was highly enriched in CoV+ inpatients (**Figure 2H).**Taken together, these data suggest SARS-CoV-2 infection is associated with a shift in the nasal microbiome and acquisition of pathobionts.

**Figure 2:**
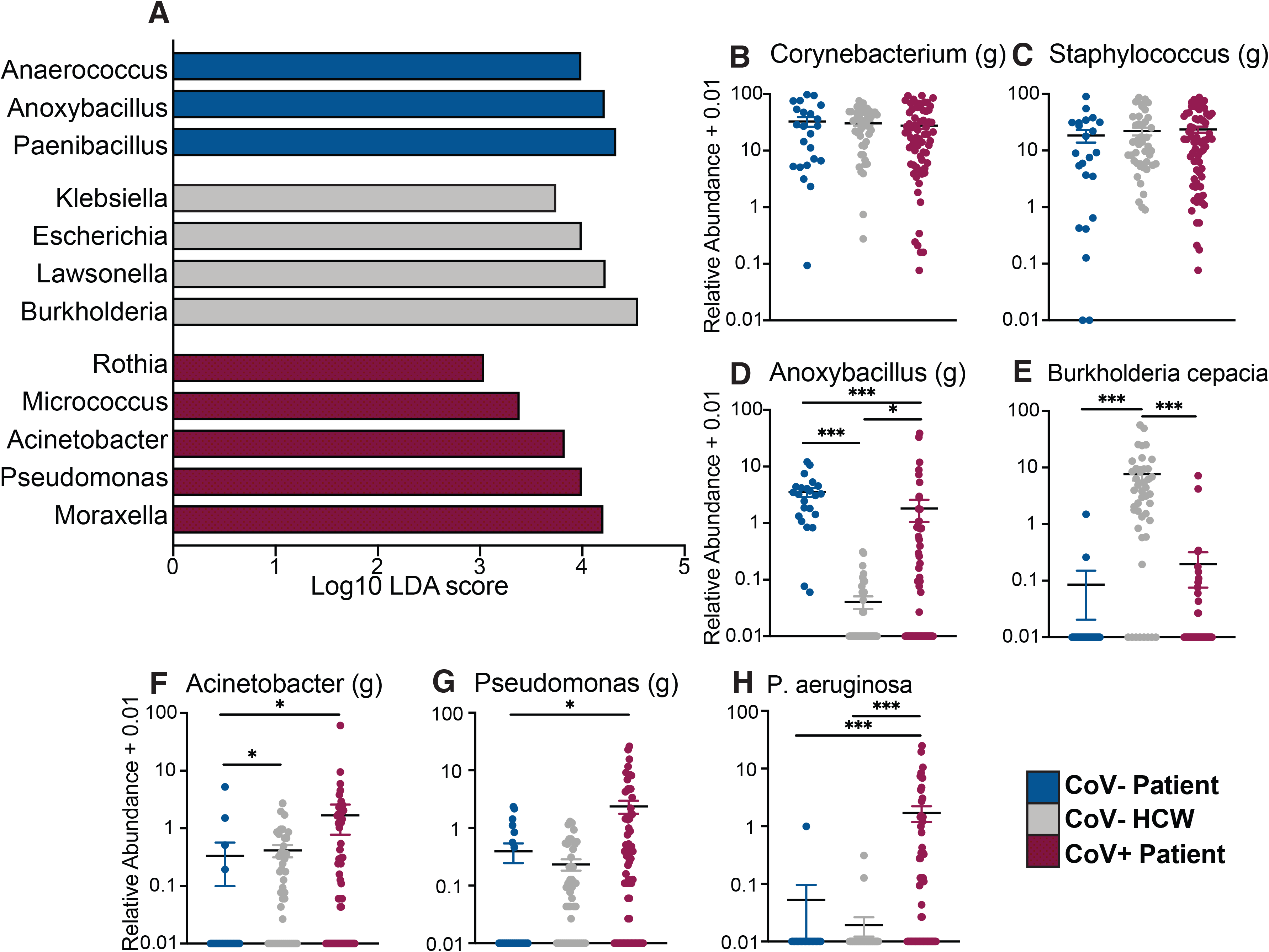
The nasal microbiome of SARS-CoV-2 infected patients and healthcare workers are enriched in opportunistic bacterial pathogens. (A) Select differentially abundant genera in the nasal microbiome between CoV-, HCW, and CoV+ individuals. Differential abundance was determined using LEFsE (Log10 LDA score > 2). (B-H) Scatter plots of bacterial genera and species of interest identified by LEFsE analysis plotted as log10 relative abundance + 0.01 and colored by host status. Horizontal black lines represent the mean of each group with whisker the SEM. Significance for panels B-H was determined using Kruskal Wallis non-parametric ANOVA, with Dunn’s multiple comparison * = p < 0.05, ** = p <0.01, *** = p < 0.001, **** = p < 0.0001.

### Associations between SARS-CoV-2 vRNA load and the nasal microbiome

We next sought to determine whether SARS-CoV-2 viral RNA (vRNA) load is associated with changes in the nasal microbiome. To this end, we divided CoV+ inpatients into three groups based on cycle threshold (Ct) values at admission: High Ct value (>34.6; n=23), Mid Ct value (32.9-25.0; n=23), and Low Ct values (<23.5; n=22) (**Supp Figure 1 D)**. We compared the number of observed ASVs, and community composition based on unweighted and weighted Unifrac distance, as well as intra-group variability across the 3 Ct groups (**Supp Figure 1 E-I**). vRNA load did not impact the diversity of the nasal microbial community (**Supp Figure 1 E-G**) However, when the relative abundance of taxa was taken into account, viral loads explained 10% of the overall community composition (**Supp Figure 1H**, PERMANOVA p value<0.001). Moreover, the nasal communities of patients with low and mid Ct groups were more similar to each other than either group was to the high Ct (**Supp Figure 1I**), suggesting that shifts in the nasal microbiome associated with SARS-CoV-2 infection may be affected by vRNA load.

We next assessed differentially abundant genera between the 3 Ct groups (**Figure 3**). LefSe analysis showed that nasal microbial communities in patients with low vRNA loads were had higher abundance of *Streptococcus*, while those of patients with moderate vRNA loads were denoted by abundance of *Corynebacterium*, and those of patients with high vRNA loads by *Cutibacterium*, *Neisseria*, and *Pseudomonas* (**Figure 3A**). The relative abundance of the genus *Streptococcus* was highest in the high CT group **(Figure 3B**), while that of *Corynbacterium* was more abundant in the Mid and Low CT groups (**Figure 3C).** Finally*, Neisseria* and *Pseudomonas* were more abundant in the low Ct group (**Figure 3D,E**) and *Pseudomonas areuginosa* was positively associated with viral loads (**Figure 3F**).

**Figure 3:**
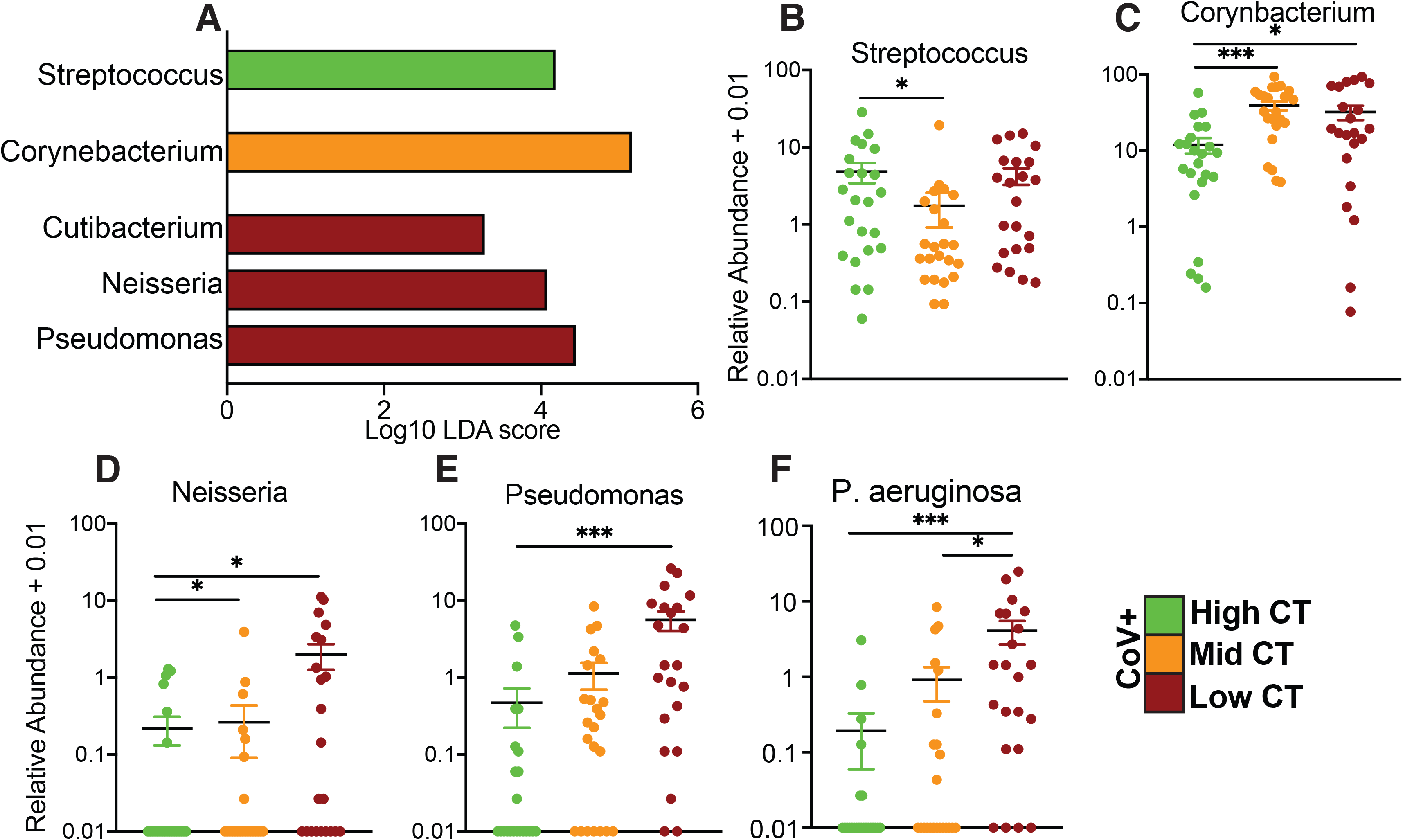
The nasal microbiome of SARS-CoV-2 infected patients is associated with viral RNA load. (A) Differentially abundant genera in the nasal microbiome between CoV+ individuals stratified by viral RNA load into High, Mid, and Low Ct. Differential abundance was determined using LEFsE (Log10 LDA score > 2). (B-F) Scatter plots of bacterial genera and species of interest identified by LEFsE analysis plotted as log10 relative abundance + 0.01 and across the three groups. Horizontal black bars represent the mean of each group with whisker the SEM. Significance for panels B-H was determined using Kruskal Wallis non-parametric ANOVA, with Dunn’s multiple comparison * = p < 0.05, ** = p <0.01, *** = p < 0.001, **** = p < 0.0001.

### SAR-CoV-2 infection remodels the nasal epithelium transcriptome

We compared the nasal transcriptomes of CoV-HCW and CoV+ inpatients using RNA-Seq to understand changes in host responses to infection (**Figure 4**). We idenitifed a total of 692 differentially expressed genes (DEGs) between the two groups (n=377 upregulated, n=315 downregulated) in the CoV+ group (**Figure 4A**). Functional enrichment showed that upregulated DEGs enriched to multiple Gene Ontology (GO) terms associated with innate and adaptive host defense pathways such as “response to virus”, and “inflammatory response”, “response to bacterium” and “lymphocyte activation” (**Figure 4B**). A large number of interferon-stimulated genes (ISGs; e.g., *HERC5*, *IFITM3, ISG15, OAS2, RSAD2*) and those related to type I interferon signaling (e.g., *IRF7, STAT1*) mapped to these GO terms “response to virus” (**Figure 4C**). Genes with roles in NFKB and JUN-ATP-1 signaling, such as *BCL3, JUN* and *MYD88*, mapped to GO terms “inflammatory response”. In terms of cell-mediated immunity, expression of genes involved in leukocyte chemotaxis (e.g., *CCL2, CCL4, CXCL9, CCR1*), hematopoiesis (e.g., *CD53, IKZF1, IKZF3*, *JAK3*) and B cell activation (e.g., IGHA, LCK, ZAP70) were upregulated in CoV+ patients. Interestingly, genes that encode inhibitory receptors (e.g.*CD300E* and *LAGLS9*) as well as genes involved in cell death (e.g. *BCL2, NUPR1* and *ZBP1*) were also highly upregulated. Additionally, upregulated DEGs (e.g. *SNCA, EGR1* and *MDK)* belonged to GO terms associated with cell death, including “neuron death” (**Figure 4B**).

**Figure 4:**
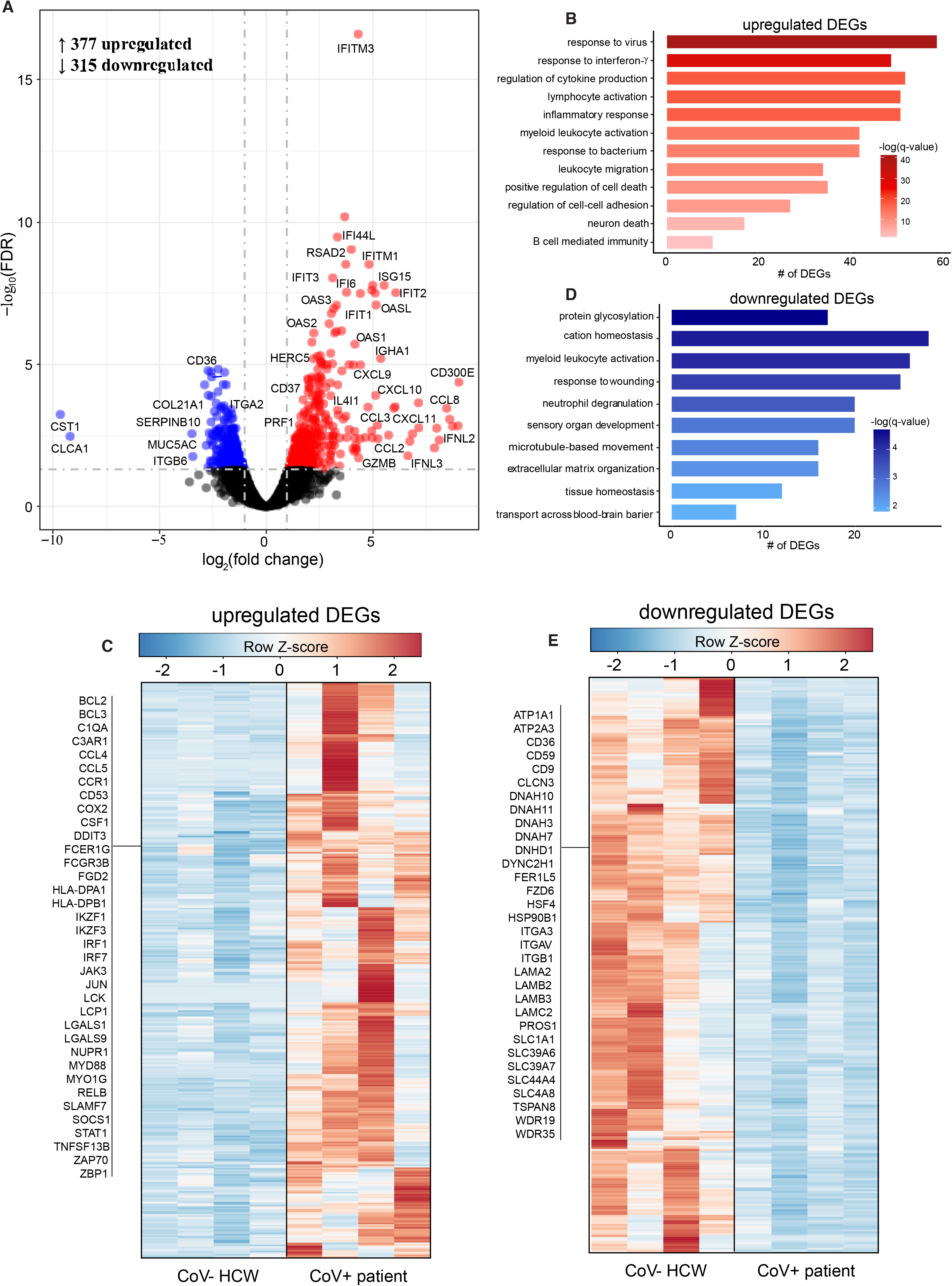
Transcriptional profiling of the nasal passages reveal robust immune activation. (A) Volcano plot of global gene expression changes in CoV+ patients relative to CoV-HCWs. Upregulated differentially expressed genes (DEGs) are indicated in red; downregulated in blue. (B) Functional enrichment of upregulated. Horizontal bars represent the number of genes enriching to each GO term with color intensity representing the negative log of the FDR-adjusted p-value [-log(q-value)]. Range of colors based on the GO terms with the lowest and highest – log(q-value) values. (C) Heatmap of upregulated. Exemplar DEGs are highlighted. Columns of all heatmaps represent the rpkm of one individual. Range of colors per each heatmap is based on scaled and centered rpkm values of the represented DEGs. Red represents upregulation; blue, downregulation. (D) Functional enrichment of downregulated DEGs as described in (C). (E) Heatmap of downregulated DEGs with exemplar DEG highlighted. See (D) for additional details

Downregulated DEGs mapped to GO terms related to maintenance of homeostasis (e.g., “response to wounding”, “), cellular organization (e.g., “microtubule-based movement”) and neuronal tissue processes (e.g., “sensory organ development”) (**Figure 4C**). Notable downregulated DEGs encoded for integrins (e.g., *ITGA2/3/V, ITGB1/6*) laminins (e.g., *LAMA2/B2/B3/C2*) and microtubules (e.g., *DNAH3/7/10/11, DYNC2H1, WDR19*) comprising intracellular and extracellular structures like cilia and cell-cell adhesion junctions. We also observed a downregulation of genes associated with mucin production in nasal passages (e.g., *MUC20, MUC5AC*). Of the ion channels downregulated (e.g., *ATP1A1, SLC39A*), several were involved in chloride ion balance and neuron homeostasis (e.g., *CLCN3, CLCA1, SLC1A1, SLC4A8*) (**Figure 4A, E**).

### Viral genome sequences do not correlate with viral loads

To identify potential relationships between vRNA load and viral evolution, we assembled and characterized SARS-CoV-2 genomes from each CoV+ in-patient (**Figure 5**). Ct value was negatively associated with viral genome coverage, with 100%, 74% and 13% of genomes from Ct groups 10-20, 21-30 and 31-40 achieving greater than 90% genome coverage, respectively (**Figure 5A**). All genomes with adequate coverage harbored the spike D614G and nsp12 P4715L mutations (**Figure 5B**). One synonymous mutation in nsp3 (F924) was also present in all genomes. The nucleoprotein T205I mutation associated with the 501.V2 variant was found in one individual (Ct 31-40). Genomes from each Ct group belonged to three main clades 20A, 20B, and 20C with no distinct relation to current variants of concern (P.1, B.1.1.7, B.1.351, B.1.427, B.1.429) or partitioning according to Ct value (**Figure 5C**). Compared to all genomes collected during 2020 in the local county (Orange County, California from January to December 2020), very few sequences belonging to the 20H clade, which emerged in October, were detected (**Fig.S3)**.

**Figure 5:**
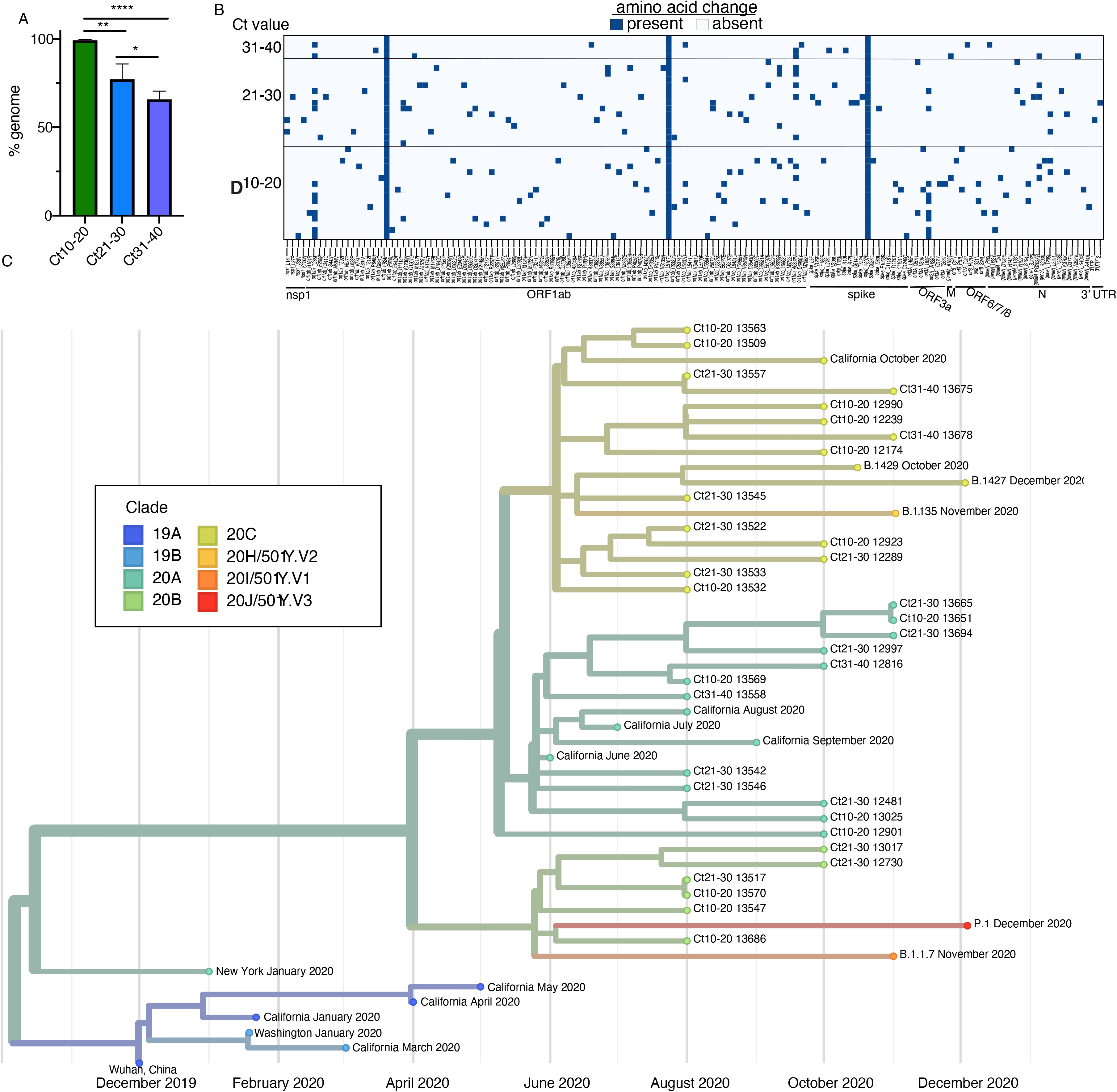
Viral RNA load and genetic variation. (A) Genome coverage (%) for each Ct group. (B) Amino acids mutations occurring in genomes with coverage > 90%. Each row represents an individual subject. Horizontal axis is order 3’ to 5’. (C) Nextstrain phylogenetic tree of genomes in (B) the original Wuhan isolate, variants of concern (B.1.1351, P.1, B.1.1.7), January 2020 Washington and New York isolates, and representative isolates from California January-September 2020 are used for comparison.

## DISCUSSION

In this study, we explored connections between acute SARS-CoV-2 infection, the nasal microbiome, and the local host transcriptional response. While a wealth of studies have focused on the systemic host response to SARS-CoV-2 and its association with disease severity, few studies have investigated the impact of acute SARS-CoV-2 on the nasal epithelium. SARS-CoV-2 infection occurs primarily via respiratory droplets with the majority of initial viral replication taking place in the nasal epithelia (*11, 14*). The majority of individual who contract SARS-CoV-2 are able to clear infection within the upper respiratory tract resulting in mild-moderate disease (*3*). However, in some individuals, the initial response is not sufficient, and SARS-CoV-2 migrates into the lower respiratory tract leading to severe COVID19 often characterized by acute respiratory distress syndrome, secondary pneumonias, and cytokine storms (*3, 4, 27, 50*). There is a clear need to understand both the host and microbial factors in the nasal cavity during acute infection that may have implications for both viral clearance and disease severity.

The nasal microbiome of a healthy individual is dominated by *Corynebacterium, Staphylococcus, Streptococcus, Dolosigranulum,* and *Moraxella (51–53)*. The composition of the nasal microbiome can impact host susceptibility and disease course following respiratory infection (*54, 55*). Additionally, acute viral infections have been shown to modulate bacterial communities potentially favoring the expansion of opportunistic pathogens (*56, 57*). In this study we show that nasal microbiome of CoV+ patients was enriched in pathogenic bacteria such *Acinetobacter Moraxella* and *Pseudomonas aeruginosa.* An expansion of *Pseudomonas* was also observed following Influenza A infection and in throat samples collected from COVID-19 patients (*58, 59*). These data suggest that expansion of *Pseudomonas* is not SARS-CoV-2 specific but rather represents a response to the inflammatory responses associated with acute viral infections. Indeed, *Pseudomonas aeruginosa* has been shown to thrive in hyperinflammatory environments such as the cystic fibrosis lung (*60, 61*). Another potential explanation, is that changes in the mucosal environment such as accumulation of mucus favor the growth of this organism. Our findings are in line with increased risk of secondary bacterial infections in COVID-19 patients, especially those who are severely ill despite high rates of antibiotic use (*62–64*). Patients with secondary infections have a poor prognosis, lower discharge rates, and higher mortality rates (*65*). Further longitudinal studies should focus on determining if these pathobionts cause secondary pneumonias associated with COVID-19. Our findings suggest that the composition of the nasal maybe provide a powerful biomarker of an individual’s risk for secondary pneumonias in viral infections.

Hospital acquired infections (HAI) are a major public health concern. A recent multi-center study found that 46% of patients hospitalized with COVID-19 suffered from a HAI (*50*). HAI are often attributed to contamination associated with healthcare workers or hospital environments (*66*). One of the major reservoirs for opportunistic pathogens is the nasal cavity which can act as the entry point and contribute to the spread of bacterial infections to other sections of the respiratory tract (*67*). It has been well established that HCW are commonly carriers of pathogens associated with nosocomial infection such as MRSA given their extended exposure to the hospital environment (*68*). Our analysis found an enrichment of pathobionts such as *Escerichia*, *Klebsiella*, and *Burkholderia cepacia* in the nasal communities of HCW. *B. cepacia* is a highly prevalent nosocomial infection in patients with underlying conditions such as cystic fibrosis (*69, 70*) and the elderly (*71*). The relative abundance of *Acinetobacter* was significantly higher in both HCW and CoV+ patients suggesting potential transfer between these two groups. Further studies should explore the role of HCW in the spread and acquisition of these pathogens in the hospital environment beyond the known pattern of MRSA.

In this study we also profiled the host transcriptional profile of the nasal epithelium in a subset of patients. Our findings highlight an upregulation of genes associated with innate immune cell activation, antiviral defense, and inflammation as recently described (*72, 73*). We also observed a transcriptional signature associated with cell death which has been proposed as a key biosignature of COVID-19 infection (*74, 75*). Furthermore, the robust upregulation of genes associated with neuronal death and downregulation of genes influencing epithelial integrity and sensory organ development supports a mechanism for infection-induced anosmia and ageusia (*24, 25, 76*). Finally, we did not detect overrepresentation of viral genomes to specific clades not well-established in California in late 2020 or partitioning by Ct value (*77, 78*). These findings suggest that viral genetic variation may have limited impact on viral replication in mucosal tissues, although studies utilizing larger cohorts are needed to confirm this hypothesis.

In summary, data presented in this manuscript show that SARS-CoV-2 infection lead to a distinct shift in the composition of the nasal microbiome including an expansion of known bacterial pathogens such as *Pseudomonas aeruginosa*. Additionally, HCW harbor nasal enriched in pathogens known to cause nosocomial infections such as *Acinetobacter* and *Burkholderia cepacia*. Our host transcriptional profiling illustrated a robust local immune response to SARS-CoV-2 infection and provided support for neuronal damage in the URT leading to anosmia. This study had some limitations. The samples used in this study were initially collected for SARS-CoV-2 diagnosis and were therefore not processed in the typical fashion for either host transcriptomics or microbiome analysis. Nevertheless, we were able to obtain enough DNA and RNA for the studies shown here. Additionally, our study consisted of only 1 time point. A longitudinal study should be performed to gain much needed insight into the dynamic changes in microbial communities and host responses within the nasal cavity. Finally, analysis of samples from individuals with asymptomatic infection would provide additional valuable insight into determinants of disease.

## Author Contributions

N.S.R., and I.M. conceived and designed the experiments. N.S.R.., A.P., A.M. B.D., A.J. and I.C. performed the experiments. N.S.R, A.P., and A.M. analyzed the data. N.S.R, A.P., and I.M. wrote the paper. All authors have read and approved the final draft of the manuscript.

## Acknowledgements

Aspects of experimental design figures were generated using graphics from Biorender.com. We wish to acknowledge the support of the Chao Family Comprehensive Cancer Center resources, award number P30CA062203 for providing the de-identified biospecimens used in this study.

---

**Supplemental Figure 1:**
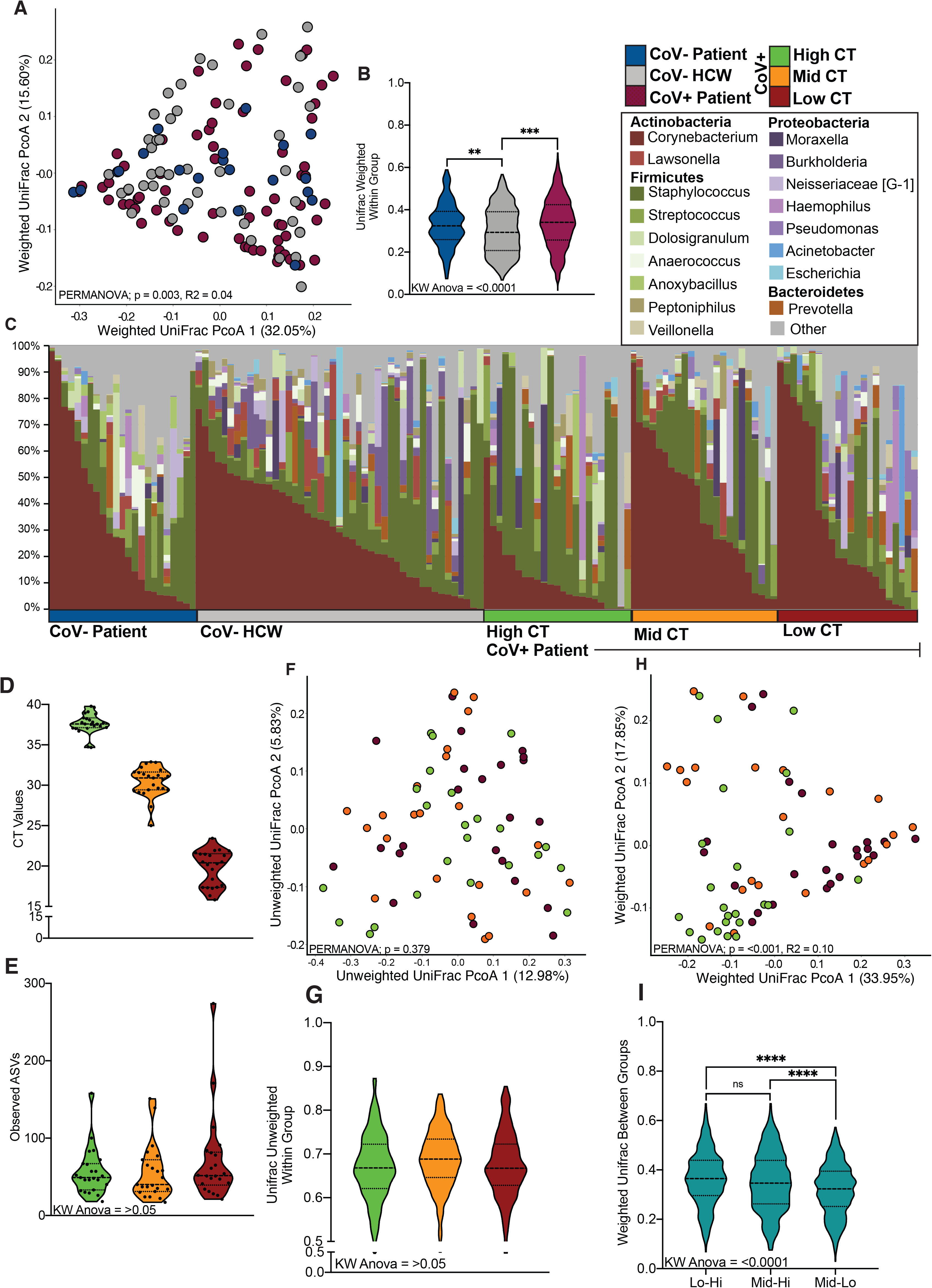
Additional features of the nasal microbiome associated with SARS-CoV-2 infection. (A) Principal-coordinate analysis of nasal microbime weighted UniFrac distance colored by host status. The contribution of host status to the total variance in the weighted UniFrac dissimilarity matrices was measured using PERMANOVA (Adonis with 10,000 permutations). (B) Violin plot illustrating average weighted UniFrac distances within each group. (C) Stacked bar of bacterial genera found at greater than 1% average abundance across the entire study population. Each vertical bar represents an individual sample and are ordered horizontally by host status. (D) Violin plot of Ct values generated via reverse transcription quantitative PCR of the SARS-CoV-2 Spike gene showing the distribution of samples within each CT group. (E) Violin plot of observed amplicon sequencing variants split by host Ct group within CoV+ patients. (F) Principal-coordinate analysis of nasal microbime unweighted UniFrac distance colored by Ct group. The contribution of Ct value to the total variance in the weighted UniFrac dissimilarity matrices was measured using PERMANOVA (Adonis with 10,000 permutations). (G) Violin plot illustrating average unweighted UniFrac distances within each group. (H) Principal-coordinate analysis of nasal microbime weighted UniFrac distance colored by Ct group. The contribution of the Ct to the total variance in the weighted UniFrac dissimilarity matrices was measured using PERMANOVA (Adonis with 10,000 permutations). (I) Violin plot illustrating average weighted UniFrac distances between each CT group. Significance for panels B,E,G,I was determined using Kruskal Wallis non-parametric ANOVA, with Dunn’s multiple comparison * = p < 0.05, ** = p <0.01, *** = p < 0.001, **** = p < 0.0001.

**Supplemental Figure 2:**
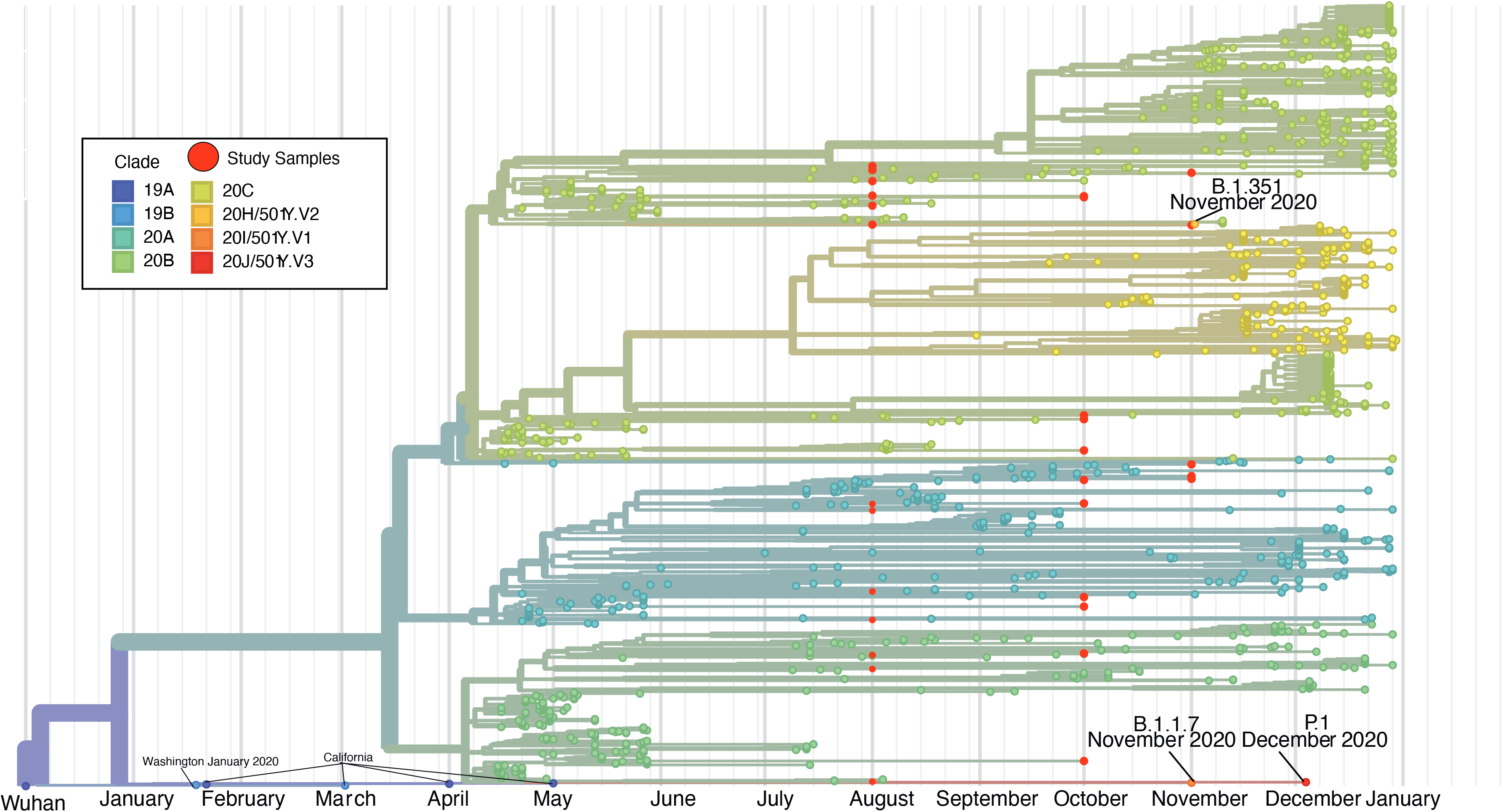
Nextstrain phylogenetic tree of genomes in Figure 5B, variants of concern (P.1, B.1.1.7, B.1.351, B.1.427, B.1.429) and all genomes collected from Orange County, California January 2020 to December 2020.

